# pyRootHair: Machine Learning Accelerated Software for High-Throughput Phenotyping of Plant Root Hair Traits

**DOI:** 10.1101/2025.07.14.664697

**Authors:** Ian Tsang, Lawrence Percival-Alwyn, Stephen Rawsthorne, James Cockram, Fiona Leigh, Jonathan A. Atkinson

**Author notes:** Joint last authorship (Fiona Leigh, Jonathan A. Atkinson).

## Abstract

Root hairs play a key role in plant nutrient and water uptake. Historically, root hair traits have been largely quantified manually. As such, this process has been laborious and low-throughput. However, given their importance for plant health and development, high-throughput quantification of root hair morphology could help underpin rapid advances in the genetic understanding of these traits. With recent increases in the accessibility and availability of artificial intelligence (AI) and machine learning techniques, the development of tools to automate plant phenotyping processes has been greatly accelerated. Here, we present pyRootHair, a high-throughput, AI-powered software application to automate root hair trait extraction from images of plant roots grown on agar plates. pyRootHair is capable of batch processing over 600 images per hour without manual input from the end user. In this study, we deploy pyRootHair on a panel of 24 diverse wheat cultivars and uncover a large, previously unresolved amount of variation in many root hair traits. We show that the overall root hair profile falls under two distinct shape categories, and that different root hair traits often correlate with each other. We also demonstrate that pyRootHair can be deployed on a range of plant species, including arabidopsis (*Arabidopsis thaliana)*, brachypodium (*Brachypodium distachyon*), medicago (*Medicago truncatula*), oat (*Avena sativa*), rice (*Oryza sativa*), teff (*Eragostis tef*) and tomato (*Solanum lycopersicum*). The application of pyRootHair enables users to rapidly screen large numbers of plant germplasm resources for variation in root hair morphology, supporting high-resolution measurements and high-throughput data analysis. This facilitates downstream investigation of the impacts of root hair genetic control and morphological variaton on plant performance.

## 3 Background

Root hairs are single cell projections that can develop on all root surfaces (Grierson et al. 2014). They play an important role in facilitating nutrient and water uptake in plants (Dolan 2017, Zhang et al. 2018) and also help to maintain root-soil cohesion (De Baets et al. 2020) and plant anchoring to soil (Haling et al. 2013). The ability of root hairs to project outwards against high soil pressure has led researchers to use root hair cells (trichoblasts) as a model system to study plant responses to mechanical resistance (Pereira et al. 2024). Despite their importance, the morphological variation and genetic control of root hairs remain relatively underexplored in comparison with other root traits (Tsang et al. 2024a). This is largely due to the inherent challenges of phenotyping root hairs, which are small in size and typically obscured by soil.

Most traditional root hair phenotyping techniques are relatively straightforward. Roots are typically grown on transparent agar plates, imaged after a few days, and the root hair measurements are manually recorded from microscope images. While this method works for small-scale studies, manual measurement of root hair traits is extremely time consuming, becomes prohibitively expensive in larger studies, and is often prone to user bias. Studies often select low numbers of root hairs to measure length at pre-defined distances from the root tip (Liu et al. 2018, Bahmani et al. 2016), and measurements are typically performed using the software FIJI (Schindelin et al. 2012, Saengwilai et al. 2021, Vatter et al. 2015, Stetter et al. 2015). Other methods involve selecting fully elongated root hairs from the mature zone of the root (Pacheco et al. 2022, Maqbool et al. 2022, Huang et al. 2020). While this method is less time consuming, it fails to provide spatial information on root hair length relative to the root tip, and constrains measurements to a specific zone of the root. Furthermore, the manual selection of the ‘longest’ root hairs per image introduces a large amount of user bias, both in terms of selection and measurement. Since the acquired image is a two-dimensional (2D) projection of a three-dimensional (3D) object, some ‘long’ root hairs may appear longer or shorter in the image than in reality. An alternative growth method is the ‘cigar roll’ technique (Liu et al. 2021, Maqbool et al. 2022), whereby seedling roots are grown on moist germination paper, and the paper subsequently rolled into a cigar shape for subsequent plant growth. While possibly faster to set up than growing on agar, this method may inflict damage to the delicate root hairs upon unrolling of the ‘cigar’ prior to imaging.

For root hair density measurements, studies often perform manual counting of individual root hairs in a defined root zone (Bahmani et al. 2016, Stetter et al. 2015). These methods are very laborious and time consuming, and are not suitable across all plant species. While the root hairs of the model dicotyledonous species *Arabidopsis thaliana* are relatively sparse and individually identifiable (Choi et al. 2019), other species such as the cereal crop bread wheat (*Triticum aestivum*) (Tsang et al. 2024b) or the nitrogen fixing model species *Medicago truncatula* (Guichard et al. 2019) have extremely dense root hairs, meaning manual counting is simply not feasible. This principle also applies to manually selecting root hairs to measure, where in species with dense root hairs, determining the start and end of individual root hairs is challenging and often impossible.

More complex methods for root hair phenotyping have also been described. For example, X-ray computed tomography (CT) has been used to model phosphate uptake in wheat root hairs (Keyes et al. 2013) and nutrient movement in the rhizosphere of rice (*Oryza sativa*) (Daly et al. 2016), providing extremely detailed characterization of root hairs in soil. Scanning and confocal microscopy techniques have also been deployed, in combination with microfludic platforms, to visualize root hairs with minimal disruption. These techniques have been used to quantify and study trichoblast nuclei at the cellular level (Aufrecht et al. 2017, Yan et al. 2021, Singh et al. 2021, Brueggeman et al. 2022). While these techniques provide a vast amount of data at extremely high resolution, the equipment required is expensive, and the methods are both computationally and labour-intensive, making them unsuitable for medium-to-large scale screens of root hair morphology.

While a number of semi-automated methods for root hair phenotyping have been described, these remain limited in the throughput that can be achieved. Vincent et al. 2017 developed a semi-automated image analysis program that quantified root hair density from images of plants grown in *in-situ* systems (e.g. rhizotrons - glass fronted soil filled chambers that allow root observation over time), which required users to manually trace outlines of their images for training. Lu et al. 2022 developed similar *in-situ* software to segment root hairs from plant roots grown in mini rhizotrons. Their method used a convolutional neural network (CNN) to perform image segmentation, enabling subsequent extraction of root hair length, diameter and area. Guichard et al. 2019 developed the software RootHairSizer, a high-throughput (42 images per hour), ImageJ-based algorithm for semi-automated measurements of root hair length, growth rate, and the differentiation zone from agar-grown images of roots. In RootHairSizer, users are required to manually define the following: the bounding regions along the root for measurement, the thresholding method and the measurement resolution. Recently, Pietrzyk et al. 2025 developed DIRT*µ*, a python-based software that utilizes machine learning to disentangle and measure individual root hairs, regardless of root hair density in the input image. However, this software is extremely computationally expensive, and potentially unsuitable for large-scale screening experiments.

Despite the advances in root hair phenotyping tools summarized above, their inability to automatically process large numbers of images with minimal manual intervention makes them potentially ill-suited to large-scale screening of root hair traits. Here, we present pyRootHair, a novel and fast software application designed for high-throughput root hair phenotyping. Primarily, pyRootHair uses a CNN for rapid and accurate image segmentation with a graphical processing unit (GPU), while also providing a simple random forest classifier (RFC) pipeline for an alternative segmentation method. For each individual image, pyRootHair extracts up to 15 summary traits and provides high spatial-temporal resolution of root hair length and area. To demonstrate its utility, we used pyRootHair to uncover significant varietal variation in all root hair traits measured across a panel of diverse wheat cultivars, and define multiple uniquely identifiable traits, including ‘root hair profile’. We further demonstrate the effectiveness of pyRootHair in processing root hair images by exhibiting its use across a range of plant species, and validate the automatically extracted traits relative to manual measurements. pyRootHair resolves a previously persistent bottleneck in root hair phenotyping. Its ability to uncover a wealth of previously undiscovered root hair phenotypic diversity across different plant species and populations, will enable future genetic and physiological studies aimed at further understanding key aspects of the ‘hidden half’ of the plant.

## 4 Data Description

The root hair dataset used, consisted of 252 wheat seedling root images. Seeds for 24 wheat cultivars listed in Table 1 were surface sterilized and germinated on agar plates. Root images were captured via light microscopy (Leica SP9) and processed using pyRootHair.

**Table 1:** Name, ploidy, common name and country of origin of all the wheat cultivars used in this study.

| Cultivar | Ploidy | Common Name | Origin |
| --- | --- | --- | --- |
| Alchemy | Hexaploid | *Triticum aestivum* (Bread Wheat) | GBR |
| Banco | Hexaploid | *Triticum aestivum* (Bread Wheat) | SWE |
| Bersee | Hexaploid | *Triticum aestivum* (Bread Wheat) | FRA/GBR |
| Brigadier | Hexaploid | *Triticum aestivum* (Bread Wheat) | GBR |
| Brompton | Hexaploid | *Triticum aestivum* (Bread Wheat) | GBR |
| Claire | Hexaploid | *Triticum aestivum* (Bread Wheat) | GBR |
| Copain | Hexaploid | *Triticum aestivum* (Bread Wheat) | FRA |
| Cordiale | Hexaploid | *Triticum aestivum* (Bread Wheat) | GBR |
| Dakter | Tetraploid | *Triticum durum* (Pasta Wheat) | FRA |
| Flamingo | Hexaploid | *Triticum aestivum* (Bread Wheat) | DNK/NLD |
| Gladiator | Hexaploid | *Triticum aestivum* (Bread Wheat) | GBR |
| Hereward | Hexaploid | *Triticum aestivum* (Bread Wheat) | GBR |
| Holdfast | Hexaploid | *Triticum aestivum* (Bread Wheat) | GBR |
| Kloka | Hexaploid | *Triticum aestivum* (Bread Wheat) | DEU/DNK/GBR |
| Kofa | Tetraploid | *Triticum durum* (Pasta Wheat) | USA |
| Maris Fundin | Hexaploid | *Triticum aestivum* (Bread Wheat) | GBR |
| Rialto | Hexaploid | *Triticum aestivum* (Bread Wheat) | GBR |
| Robigus | Hexaploid | *Triticum aestivum* (Bread Wheat) | GBR |
| Slejpner | Hexaploid | *Triticum aestivum* (Bread Wheat) | DNK/SWE |
| Soissons | Hexaploid | *Triticum aestivum* (Bread Wheat) | FRA |
| Spark | Hexaploid | *Triticum aestivum* (Bread Wheat) | GBR |
| Steadfast | Hexaploid | *Triticum aestivum* (Bread Wheat) | GBR |
| Stetson | Hexaploid | *Triticum aestivum* (Bread Wheat) | GBR |
| Xi19 | Hexaploid | *Triticum aestivum* (Bread Wheat) | GBR |

## 5 Analyses

### 5.1 Variation in Root Hair Traits

When using pyRootHair to extract root hair phenotypic data from images across a panel of 24 wheat cultivars, a large amount of variation was observed for all root hair traits extracted (Figure 3). Mean root hair length (RHL) varied between 1-2 mm for most cultivars. Within the bread wheat cultivars, Bersee had notably shorter root hairs relative to all other accessions (Figure 3A). The durum wheat varieties Dakter and Kofa had shorter average RHL than most bread wheat genotypes. Mean RHL exhibited a strong positive correlation with maximum RHL (Figure 4).

**Figure 1:**
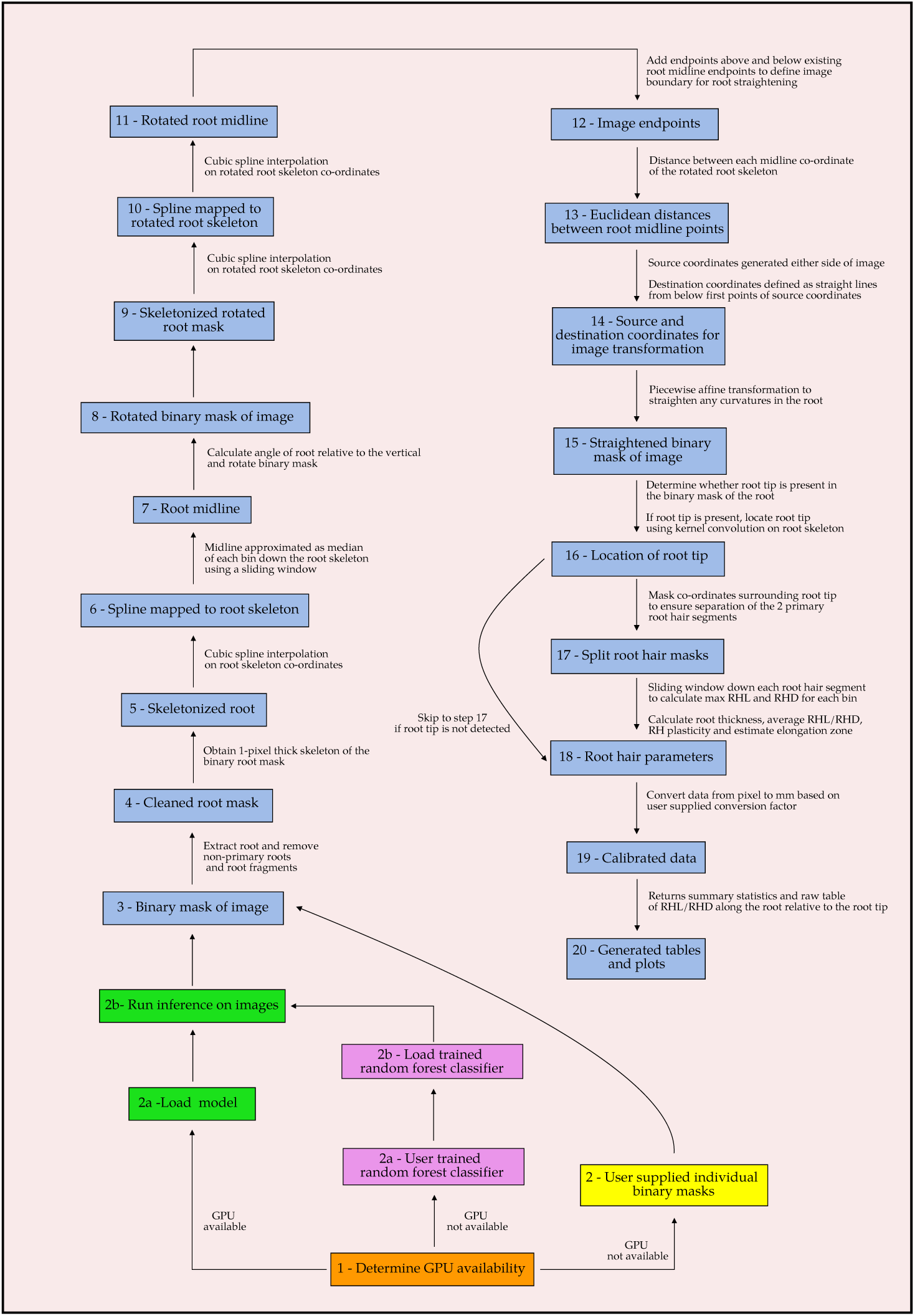
pyRootHair pipeline methodology overview. Blue boxes illustrate primary input/output of each image processing step. Green boxes illustrate workflow steps for the default pipeline using a graphical processing unit (GPU). Red boxes illustrate steps taken for the alternative pipeline (for users without access to a GPU) using a random forest classifier (RFC) model. The yellow box illustrates a step for users supplying individual binary masks for data. RHA = root hair area, RHL = root hair length.

**Figure 2:**
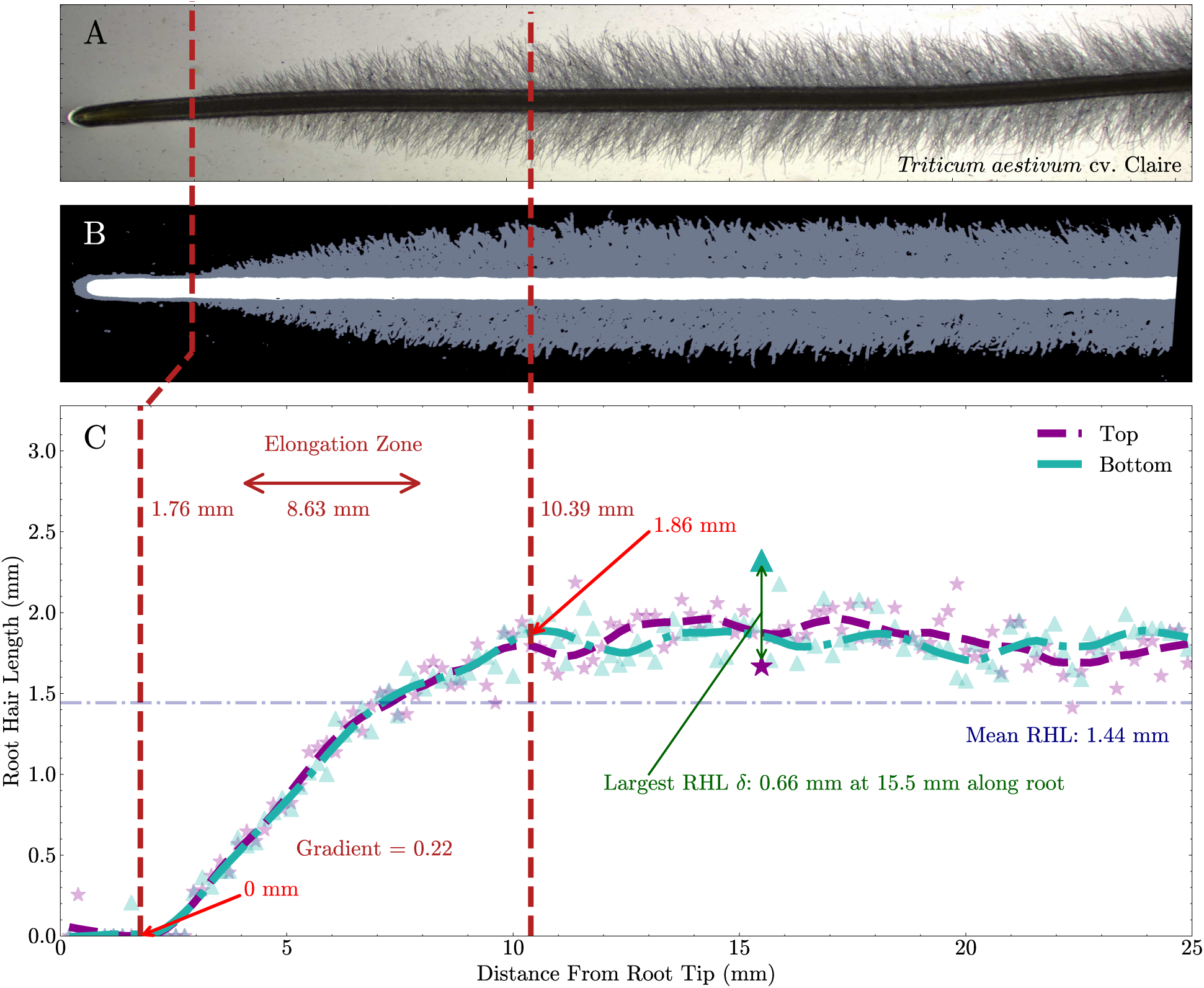
Visual representation of selected root hair traits extracted from an example bread wheat (cv. Claire) image using pyRootHair. A) Raw image. B) Binary mask from segmentation of the raw image. C) Green and purple markers illustrate root hair lengths along the root. Length profile for each segment (top or bottom) is illustrated by dashed regression lines through the markers. Vertical dashed maroon lines indicate the automatically identified root hair elongation zone, calculated as the largest region of continuous upward trajectory in the root hair profile. Gradient of the elongation zone is calculated as the maximum change in root hair length within the zone, divided by the length of the zone. Mean root hair length is calculated across the entire root length, illustrated by the dashed horizontal orange line. The maximum difference (*δ*) in root hair length between the root hair segments is 0.66 mm at 15.5 mm away from the root tip.

**Figure 3:**
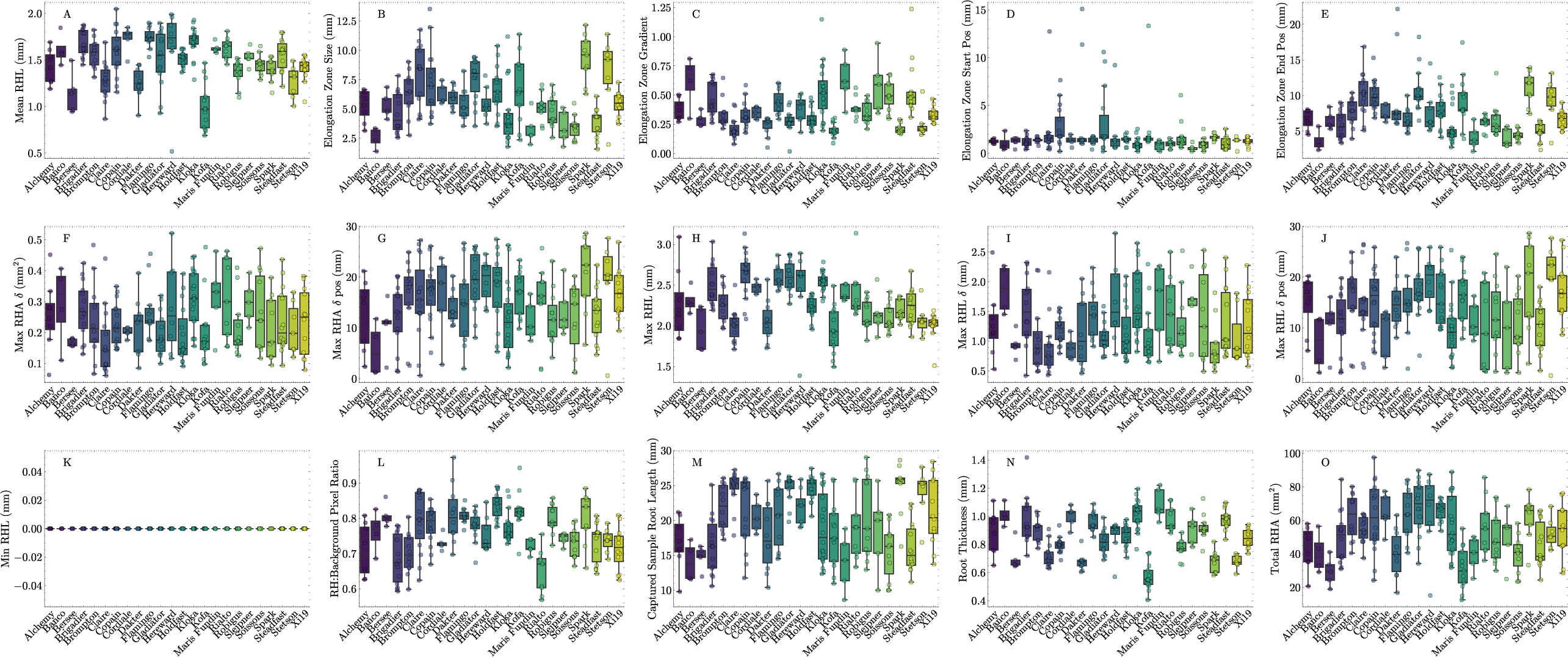
Diversity in root hair traits across all screened wheat cultivars. Boxplots illustrate spread of data for each cultivar and each trait. The horizontal lines within each boxplot illustrates the median, whiskers represent the data range, and data from an individual image is displayed as a dot.

**Figure 4:**
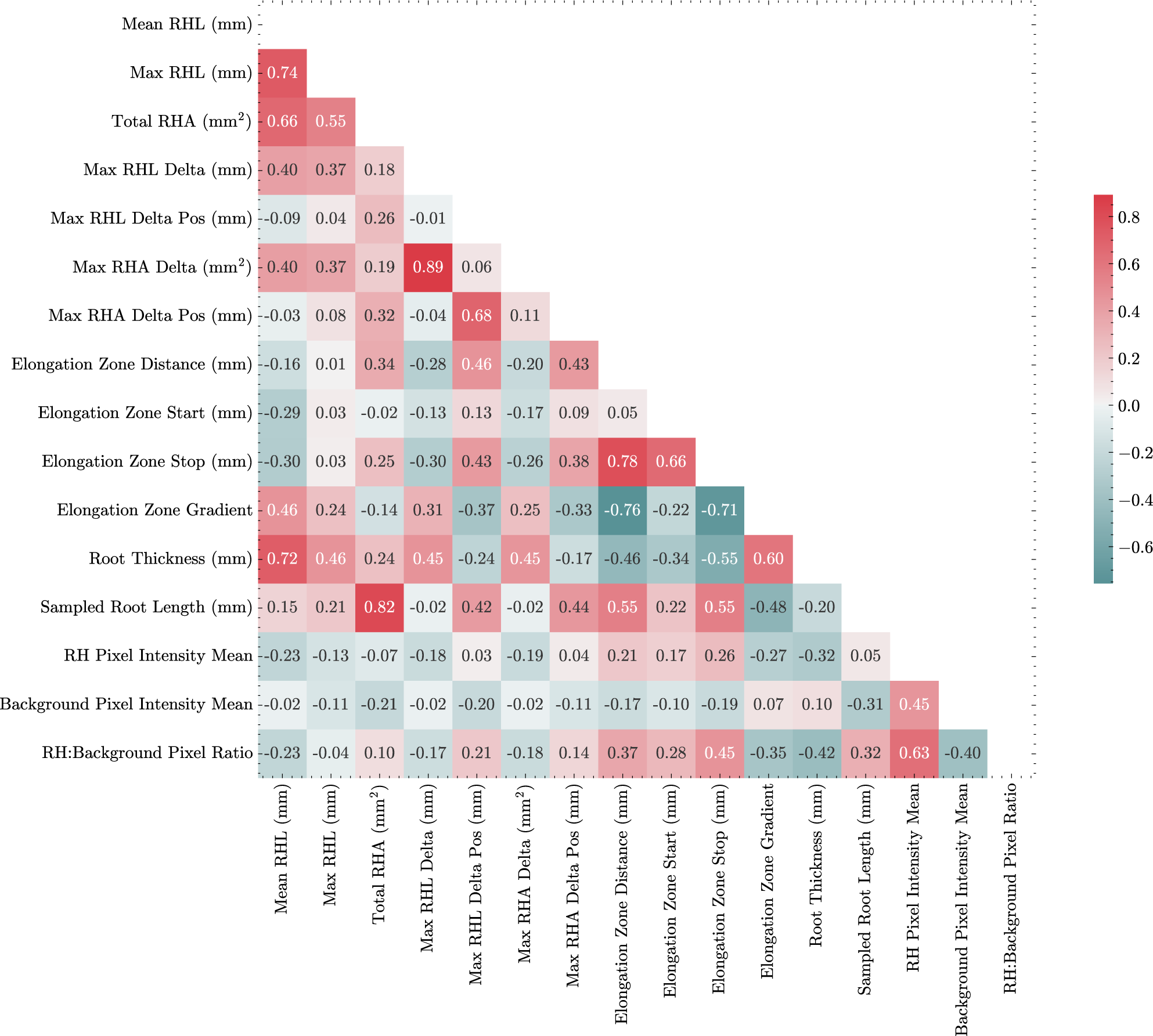
Correlation matrix illustrating relationships between root hair traits measured across the 24 wheat cultivars using pyRootHair. Numbers within squares illustrate Pearson correlation coefficients.

Root hair area (RHA) was calculated as the total area occupied by the root hairs in each binary mask. Total RHA (Figure 3O) was positively correlated with mean RHL (Figure 4, *R*^2^ = 0.66). This was unsurprising, as RHA was simply measured as the 2D area occupied by the root hair mask. As such, images that contained a longer section of root would have a higher mean RHL and thus a higher RHA due to the presence of more root hairs. Root hair density (RHD) was estimated using the RH:Background Pixel ratio (Figure 3L). This ratio was calculated as the mean pixel intensity of the root hairs in the input image, divided by the mean pixel intensity of the background. In backlit images, the lighter background would have a higher mean pixel intensity, while the root hairs would be darker due to their obstruction of light and thus have a lower mean pixel intensity. If a cultivar had more root hairs per unit area of root, the root hairs would be darker, decreasing the pixel intensity value of the root hairs. Thus, assuming consistent lighting across input images, lower RH:Background ratios could indicate a cultivar had higher RHD, while higher RH:Background ratios could indicate a cultivar had lower RHD (Supplementary 2B). Furthermore, the variation in mean background pixel intensity within each cultivar illustrates the consistency of lighting conditions between images, and can therefore serve as a quality control metric.

Here, the trait ‘root hair plasticity’ was determined as the variation in root hair length and area within an individual root image (Figure 2C). Bread wheat cultivars, including Brigadier, Brompton, Hereward and Soissons, all exhibited large fluctuations in max RHL *δ* between the left and right root hair segments (Figure 3I). The position along the root of maximum RHL fluctuation also varied between cultivars (Figure 3J), but no linear relationship with Max RHL *δ* was found. In contrast, some cultivars (notably Robigus, Spark and Steadfast) exhibited minimal variation in RHL *δ* between individuals, indicating these cultivars had more uniform root hair growth.

The elongation zone was defined here as the largest continuous region where root hairs experienced positive growth (Figure 2C, region bound by dashed red lines). Notably, the bread wheat cultivar Spark had a very large elongation zone size (Figure 3B). Three cultivars (Claire, Copain and Kloka) all exhibited wide variation in elongation zone size between biological replicates. The starting position of the elongation zone was relatively consistent between all cultivars (Figure 3D). In contrast, the end position of the elongation zone varied more relative to the start position (Figure 3E). The gradient of the elongation zone was negatively correlated with the stop position (Figure 4, *R*^2^ = *−*0.71) and size (Figure 4, *R*^2^ = *−*0.76).

All cultivars were imaged five days after germination. Across all cultivars, variation in root length between individuals were observed. This was likely due to underlying genetic differences, plate and positional growth room effects. As such, each image captured was positioned to contain the maximum length of root suitable for downstream processing. Thus, for cultivars with a consistent level of root growth (e.g. Gladiator, Spark, Stetson), there was minimal variation in captured sample root length. Conversely, cultivars with a high degree of variation in root length (e.g. Brigadier, Copain, Kloka) displayed large variation in captured sample root length (Figure 3M). The thickness of the captured root (i.e mean width of the root) (Figure 3M) varied across all cultivars. Unsurprisingly, no linear relationship was found between captured root length and root thickness. Root thickness was positively correlated with mean RHL (Figure 4, *R*^2^ = 0.72), elongation zone gradient (Figure 4, *R*^2^ = 0.59), and negatively correlated with the elongation zone stop position (Figure 4, *R*^2^ = *−*0.54).

The greatest variation in overall root hair profile was found around the root tip, where root hairs had begun to emerge. For each cultivar, the mean root hair profile was calculated as the mean RHL of all replicates for each position along the root (Figure 5A).

**Figure 5:**
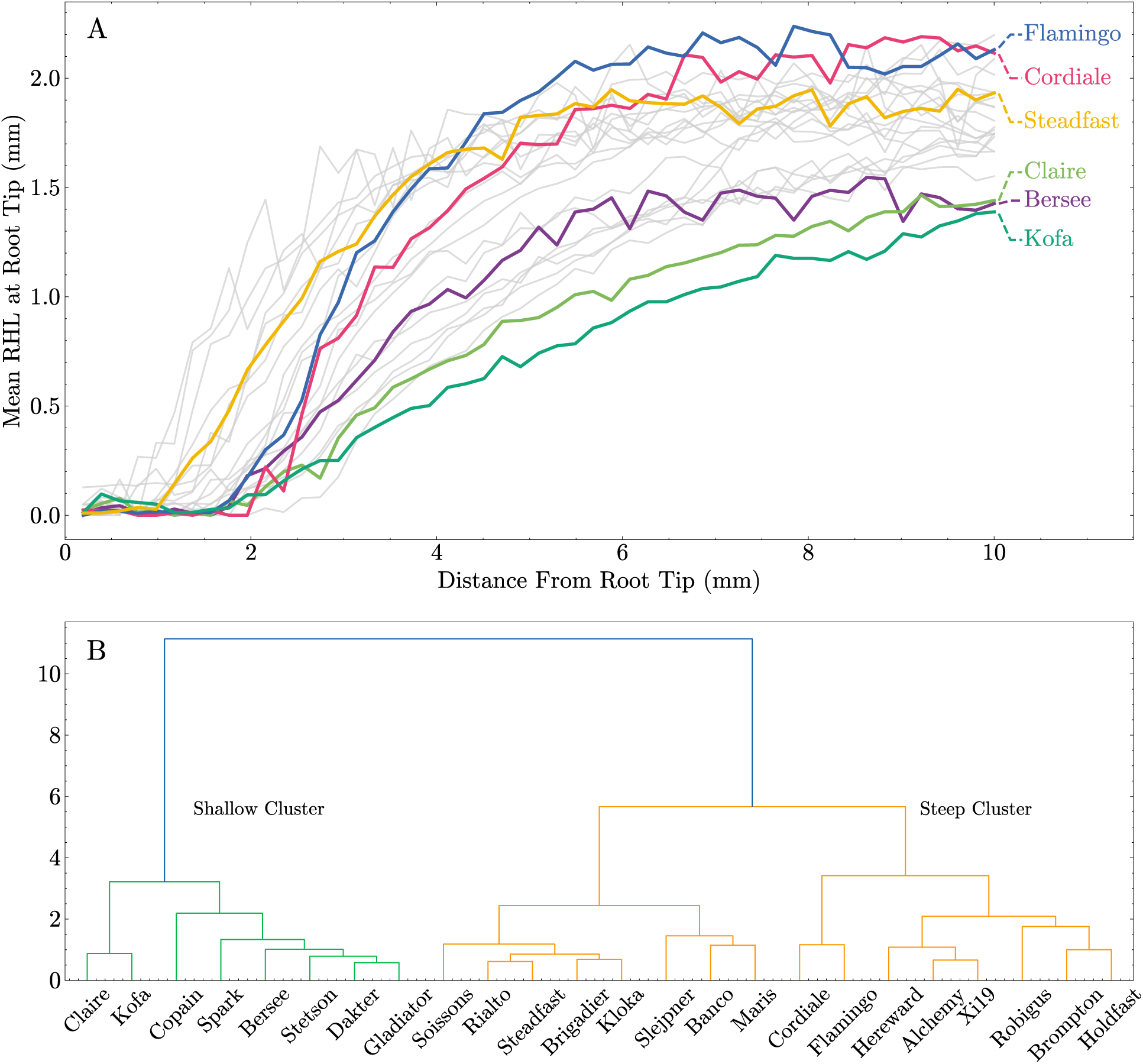
Variation in root hair profile around the root tip (1cm) across the 24 wheat cultivars phenotyped using pyRootHair. A) Mean root hair length (RHL) profile, highlighting cultivars exhibiting extreme variation in RHL within the region displayed. B) Unsupervised agglomerative clustering of root hair profiles highlights two main clusters of root hair profiles around the root tip: ‘Shallow’ (Green cluster) and ‘Steep (Orange cluster)’. See Supplementary Figure 2A for examples of cultivars with ‘shallow’ and ‘steep’ profiles.

A large degree of variation was found for both root hair emergence (minimum distance from root tip where hairs begin emerging) and overall root hair profile across the 1 cm section from the root tip (Figure 5A). Unsupervised clustering of the root hair profiles revealed two major clusters, termed here ‘Shallow’ and ‘Steep’ (Figure 5B). Cultivars in the ‘Shallow’ cluster (including Kofa, Bersee, Claire) had a shallower root hair profile gradient (Figure 3C, 5A, Supplementary Figure 2A). In contrast, cultivars in the ‘Steep’ cluster (including Flamingo, Cordiale, Steadfast) had a steeper root hair gradient, and generally had longer root hairs around the root tip compared to cultivars in the ‘Shallow’ cluster (Figure 5A, Supplementary Figure 2A).

### 5.2 Adaptability

Given that overall root morphology is relatively consistent throughout many higher plant species, the methodology deployed in the pyRootHair workflow can be translated to a wide range of wild and cultivated plant species. Here, we demonstrated that pyRootHair can accurately segment and quantify root hair traits by detailed investigation of wheat, as well as further investigation in single accessions from seven other species: oat, rice, teff, tomato (Figure 6), as well as medicago, brachypodium and arabidopsis.

**Figure 6:**
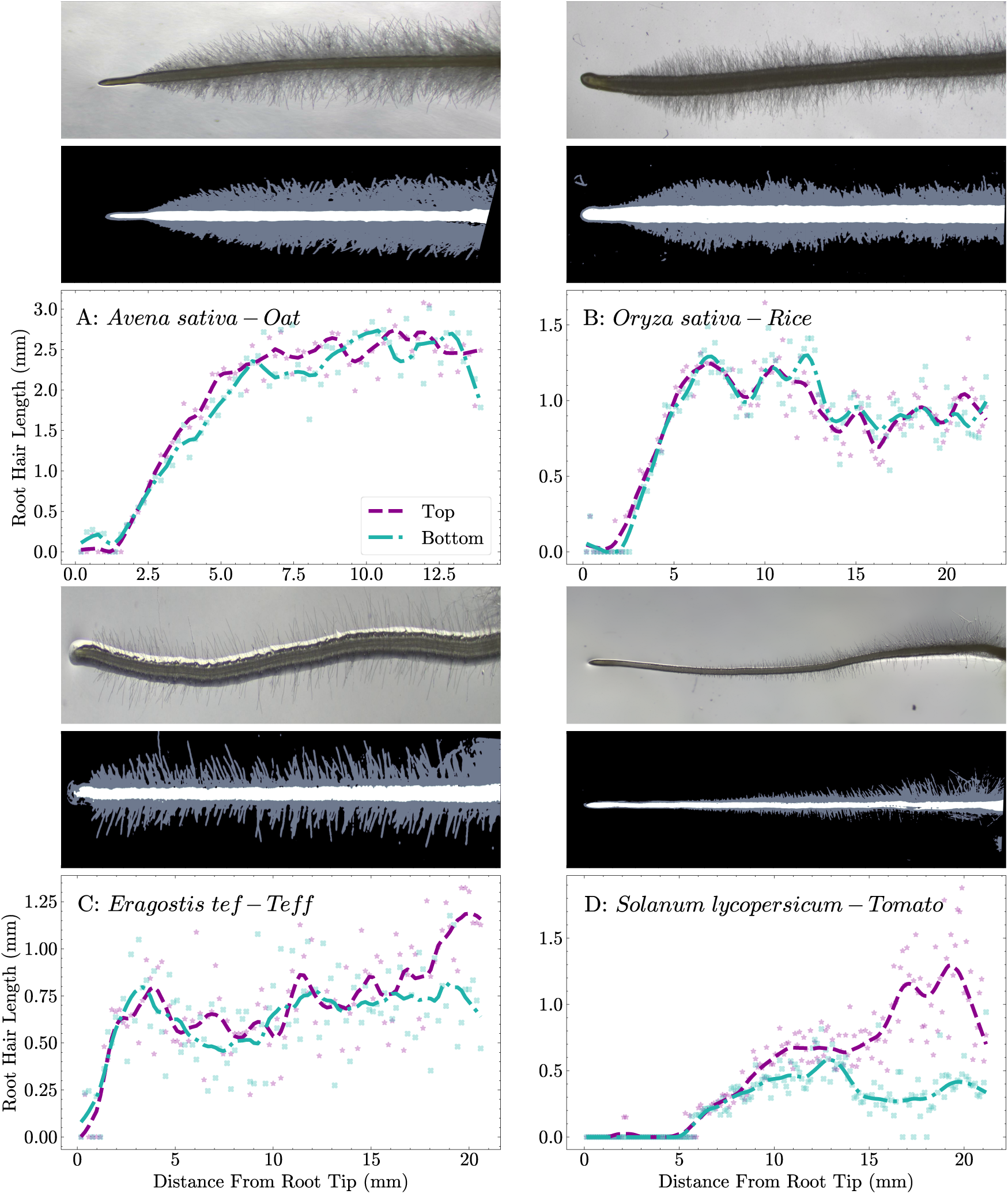
pyRootHair deployed on root images captured from different plant species. A) oat, B) rice, C) teff, D) tomato. For each panel: raw images on top, predicted binary masks in the middle, automated plots of root hair length (RHL) on the bottom. Raw images and binary masks not to scale. Top = top segment of root hairs, Bottom = bottom segment of root hairs.

### 5.3 Trait Validation

To validate traits extracted by pyRootHair, the RHL, elongation zone length and root length were manually measured across different images. Manual RHL measurements were taken from five different images, with measurements performed at 0.1 mm intervals along the root across both root hair segments. Manual measurements of the elongation zone and root length were taken from 54 and 68 different images respectively. All manual measurements were calibrated using the conversion factor of 102 pixels per 1 mm.

Strong positive correlations between automated and manual measurements were found for RHL (Figure 7A, *R*^2^ = 0.88), elongation zone length (Figure 7B, *R*^2^ = 0.55) and root length (Figure 7C, *R*^2^ = 0.97). The CNN model exhibited accurate segmentation of input images compared to manually annotated binary masks, with intersection over union (IoU) scores *>* 0.8 across different example images (Supplementary Figure 3).

**Figure 7:**
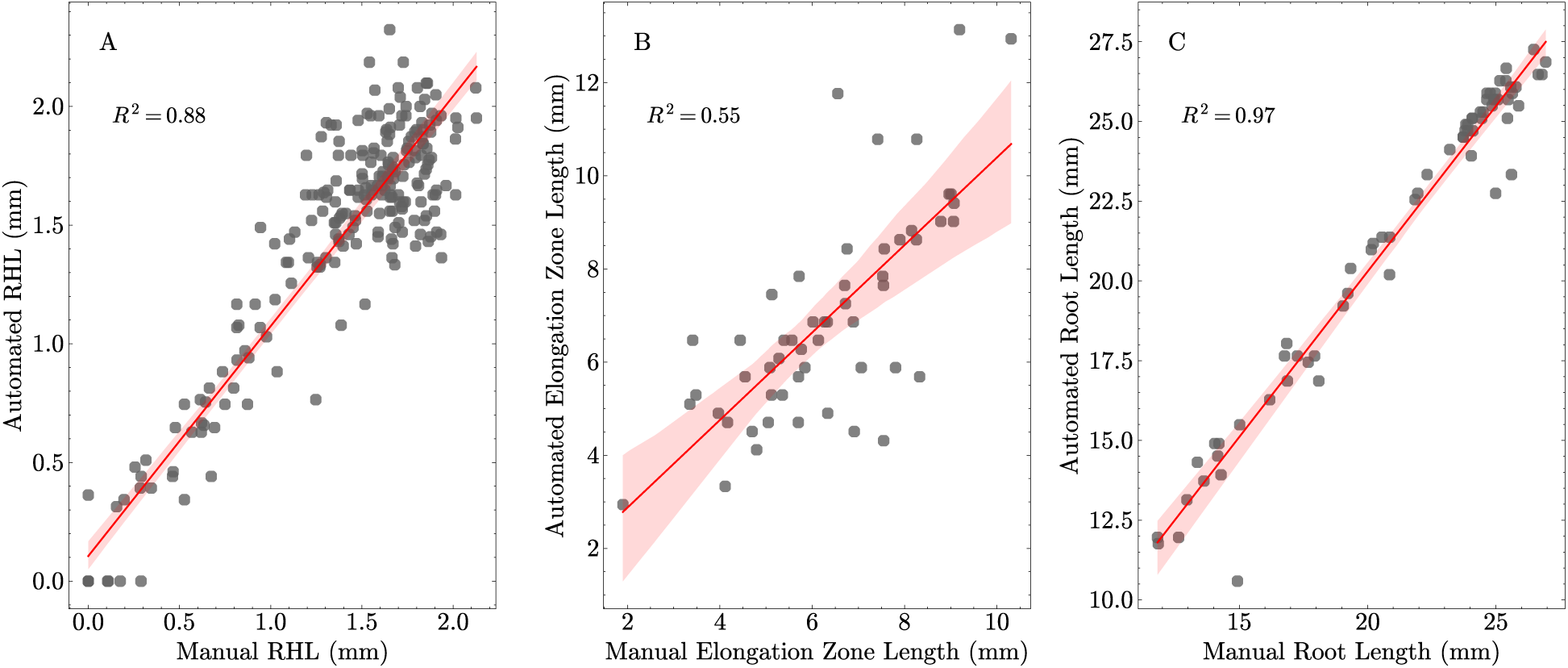
Correlation (*R*^2^) between manually measured traits and automated trait extraction using pyRootHair. A) Root Hair Length (RHL). B) Elongation Zone Length. C) Root Length. Regression line with 99% confidence interval illustrated in red.

### 5.4 Performance

Across all images for the 24 wheat cultivars, the mean per-image processing time (including inference) was 7.82 s with an Nvidia L40S GPU with 8 Gigabytes (GB) of Video Random Access Memory (RAM), compared to 7.97 s per image using the RFC pipeline (Supplementary Figure 1A) on a compute node with 20 GB RAM. The mean processing time per image was 430 s when using the CNN to perform inference without a GPU (Supplementary Figure 1A), which was significantly slower than either the GPU or RFC pipelines (Supplementary Figure 1B). No significant difference in mean image processing time was found between the two pipeline configurations. For this comparison, the RFC was trained on a single input image, and used to perform inference on all subsequent images. As such, the small sample size resulted in less accurate segmentation of input images compared to the CNN deployed in the main pipeline. Despite the reduction in segmentation accuracy with the RFC model, the extracted summary data for all wheat images displayed relatively linear relationship with the summary data from the main pipeline with a GPU (Supplementary Figure 1C, D)

## 6 Discussion

Advances in computer vision and AI have greatly accelerated the field of plant phenotyping. Here, we present pyRootHair, a rapid and new computer vision-based software application for high-throughput extraction of root hair traits from root images. To our knowledge, pyRootHair extracts more traits per input image than any other available software. Furthermore, unlike previously developed software, pyRootHair is extremely quick and enables large scale batch processing of images. Importantly, no user input is required during computation. As such, pyRootHair significantly improves the speed and accuracy of root hair phenotyping. The throughput achievable with pyRootHair means that the bottleneck in acquiring root hair data has transitioned from the phenotyping step to the plant growth and image acquisition process.

The limitations of pyRootHair primarily relate to the curvature of root hairs, and the segmentation performance. While straightening of the input root is effective at standardising the measurement along each root hair segment, RHL is calculated as the width of the root hair binary mask, and does not account for the inherent curvature within the individual root hairs. Resolving the length of individual hairs is computationally expensive and has already been accomplished (Pietrzyk et al. 2025). Rather, pyRootHair attempts to quantify overall root hair morphology of an image rather than individual hairs, which we propose is more biologically relevant. This approach is faster, and avoids any user selection biases towards longer root hairs. Secondly, accuracy and effectiveness of trait extraction is dependent on the segmentation performance. While the CNN deployed by the main pipeline is extremely flexible and powerful, training instances are laborious to annotate, and cannot be generated to cover all examples of end user images. The RFC pipeline is computationally inexpensive, but far less flexible compared to the main pipeline with the CNN.

The variation in root hair morphology identified across the 24 wheat cultivars investigated here highlights useful entry points for future genetic and physiological investigation of root hair morphology on plant performance. In particular, traits such as the shape of the root hair profile and plasticity have, to our knowledge, not been previously quantified. Identification and formalization of these new traits increases the likelihood of identifying genetic loci and genes controlling root hair morphology via forward genetic analyses, and for their physiological investigation. For example, given that root hair length is positively correlated with phosphorous uptake in different crops (Gahoonia et al. 2004, Saengwilai et al. 2021), automated RHL calculation via pyRootHair serves as an extremely convenient and fast method of pre-screening a panel of cultivars for field trials or breeder selection.

## 7 Potential Implications

Due to the throughput achieved and ease of use, we believe that pyRootHair can significantly reduce the bottleneck in root hair phenotyping, and serve as a key tool for uncovering the genetic loci and genes controlling root hair morphology in crops and other plant species. Through this process, candidate genes controlling root hair traits can be identified and selectively bred into existing populations to develop more resilient, nutrient-efficient crops for the future.

## 8 Methods

### 8.1 Plant Material

All plant material is summarized in Table 1. The 24 wheat cultivars were sourced from field-grown trials in Cambridgeshire. The accessions included 22 bread wheat cultivars, encompassing all founder lines within the ‘Niab Elite MAGIC’ (Multi-Parent Advanced Generation Inter-Cross) and ‘Niab Diverse MAGIC’ populations (Mackay et al. 2014, Scott et al. 2021). Two durum wheat (*Triticum turgidum* ssp. *durum*) varieties (Dakter, Kofa) were also included. Seeds for oat, rice, teff and tomato were either sourced from locally available seed stocks at Niab or commercially purchased. Medicago, brachypodium and arabidopsis images were provided by Guichard et al. 2019.

### 8.2 Image Acquisition

Cold-treated seeds (stored for five days at 4*^◦^*C) were surface sterilized with 70% ethanol for 60 s, then immersed in 20% bleach (sodium hypochlorite) for 10 minutes. Seeds were then washed three times with sterile distilled water in a laminar flow hood, and plated onto square agar plates (Sigma-Aldrich, 120 x 120 x 17 mm) with sterile forceps. Four seeds were placed on each plate, with each seed positioned in a corner with the embryos oriented towards the centre of the plate. Each agar plate contained 50 mL of media prepared with of 0.5 g sucrose (Sigma-Aldrich), 0.15 g phytagel (Sigma-Aldrich) and 0.2 g Murashige and Skoog basal salt (Sigma-Aldrich), mixed with 50 mL dH_2_O. Plates were sealed with micropore tape (3M) and placed horizontally on flat racks in a well lit, 25*^◦^*C growth room for five days under constant lighting. Subsequently, the roots were imaged using a Leica S9D Stereomicroscope at 0.6*×* magnification.

### 8.3 Segmentation Model

To generate and train the segmentation model, we used the self-configuring CNN generator nnU-Netv2 (Isensee et al. 2021). Binary masks of training instances were manually curated using the interactive segmentation tool ilastik (Berg et al. 2019). For each binary mask, pixels associated with the background were labelled with a 0, root hairs as 1, and roots as 2. Training instances varied in input dimensions, clarity and lighting conditions. The training set was composed of 44 wheat, 5 arabidopsis, 14 medicago, 3 brachypodium, 4 teff, 4 maize (*Zea mays*) and 9 rice images, along with the corresponding binary masks.

Model training was carried out on the Crop Diversity High Performance Computer (HPC) Cluster (Percival-Alwyn et al. 2024) using an NVIDIA A100 SXM4 80GB Tensor Core GPU. The ‘nnUNetResEncUNetMPlans’ planner preset was used for all nnU-Net stages (Isensee et al. 2024). Training was conducted with the ‘all’ fold and the ‘2d’ configuration arguments. Image augmentation was automatically performed by the nnU-Net pipeline. Z-score normalization was carried out for the three input image channels (red, green, blue). The model was trained for 1000 epochs with a batch size of 13. The CNN architecture comprised 7 convolutional layers with an increasing number of feature maps: 32, 64, 128, 256, 512, 512, 512, each with a kernel size of 3. A stride length of 2 was used for all layers except the first, which used a stride length of 1. Leaky ReLU was used as the activation function. The mean validation dice score of the model was 98% across all training instances.

### 8.4 Pipeline Configurations

To accommodate a variety of end-user hardware, pyRootHair was designed to operate across different levels of computing power, including HPC systems and personal computers. As such, different pipeline configurations are available for the end-user.

By default, the main pipeline utilizes a GPU to perform inference on input images. GPU requirements vary depending on input image sizes (Figure 1, Steps 2a,b, green). For reference, an Nvidia L40S GPU with 8 GB VRAM was sufficient to perform inference on input images of size 2600 x 1500 x 3. Inference without a GPU is still possible on a CPU, but speed will be significantly affected.

For users without a GPU, pyRootHair offers a simple alternative pipeline. Users can easily train an RFC segmentation model on a representative image of their choice with a single command. The trained RFC model can be subsequently used to run inference on input images (Figure 1, Steps 2a,2b, purple). Alternatively, end users can generate their own binary masks, which can then be individually processed for trait extraction without a GPU (Figure 1, Step 2, yellow).

### 8.5 Workflow

For a given batch of input images with the main or RFC segmentation pipeline, the predicted masks are generated first. The post-processing steps of each binary mask obeys the following methodology for trait extraction:

For each mask, the root is first extracted and segmentation noise removed (Figure 1, Step 4). The root is then skeletonised to a single-pixel wide representation of the root (Figure 1, Step 5). A spline (smooth curve fitted through a set of points) is subsequently mapped to the skeleton co-ordinates, and the root midline is approximated via computing the median of each bin in a sliding window down the root mask (Figure 1, Steps 6 and 7).

To standardise trait extraction, roots must be oriented downwards in the input image. The orientation of the root in the binary mask is calculated from the approximated root midline, and rotated such that the root tip points downwards (Figure 1, Step 8). The rotated root mask is re-skeletonised and co-ordinates are re-mapped to approximate the midline of the rotated root (Figure 1, Step 11). Next, to minimize the effect of any inherent root curvature within the rotated root, the entire binary mask is straightened via affine transformation, producing a completely straightened binary mask of the original image (Figure 1, Steps 12-15).

After straightening the mask, the root tip is located using kernel convolution along the skeleton of the straightened root (Figure 1, Step 16). The binary mask of the root hairs are then split around the root tip, separating the total root hair mask into an individual root hair mask for each side of the root (Figure 1, Step 17).

To extract RHL, the longest RHL is calculated in each bin from a sliding window along each root hair segment, starting at the root tip. The size of the window can be controlled via the --resolution argument. The default value of 20 pixels per bin size was used to compute traits from all images analysed in this paper. RHA is calculated as the total pixel area of root hair mask within each bin. The root hair elongation zone is defined as the region where root hairs experience continuous positive growth. The ‘profile’ of the root hair segment is modeled using a LOWESS (locally weighted scatterplot smoothing) regression line, and the elongation zone position, size and gradient are extracted from the longest region of the root where the gradient of the root hair profile continuously increases (Figure 2C). The plasticity of root hair growth is defined as the difference in RHL and RHA between the ‘left’ and ‘right’ sides of the root hair segment. The greatest difference in RHL and RHA between each root hair segment are recorded as ‘Max RHL/RHA *δ*’, along with the corresponding position along the root (Figure 2C). All traits are translated from pixel measurements to mm measurements, where the pixel:mm conversion factor can be controlled via the --conv argument. A full list of extracted traits is available in Table 2.

**Table 2:** Summary traits calculated for each input image. RHA = root hair area, RHL = root hair length.

| Trait | Explanation |
| --- | --- |
| Name | Name of image file |
| Batch ID | Batch ID specified via -b/--batch_id |
| Avg RHL (mm) | Average root hair length of the input image, calculated across the entire length of the root |
| Max RHL (mm) | Maximum recorded root hair length in the input image |
| Min RHL (mm) | Minimum recorded root hair length in the input image |
| Total RHA (mm<sup>2</sup>) | Total root hair area of the input image |
| Max RHL Delta (mm) | Largest difference in root hair length between each root hair segment on either side of the root |
| Max RHL Delta Pos (mm) | Position along the root corresponding to the largest difference in root hair length |
| Max RHA Delta (mm) | Largest difference in root hair area between each root hair segment on either side of the root |
| Max RHA Delta Pos (mm) | Position along the root corresponding to the largest difference in root hair area |
| Elongation Zone Distance (mm) | Length of the elongation zone, defined as the largest continuous region where root hairs are emerging |
| Elongation Zone Start (mm) | The position along the root where the elongation zone starts, calculated as distance from the root tip |
| Elongation Zone Stop (mm) | The position along the root where the elongation zone ends, calculated as distance from the root tip |
| Elongation Zone Gradient | The gradient of the elongation zone, calculated as $\delta$ RHL in elongation zone/length of elongation zone |
| Root Thickness (mm) | Average thickness of the root, calculated via a sliding window down the binary root mask |
| Captured Sample Root Length (mm) | Total length of the straightened root captured within the input image |
| RH Pixel Intensity Mean | Mean pixel intensity of the area occupied by the root hairs in the input image |
| Background Pixel Intensity Mean | Mean pixel intensity of the background in the input image |
| RH: Background Pixel Ratio | RH pixel intensity divided by background pixel intensity |

### 8.6 Output

For a given batch of input images, pyRootHair produces a summary table and a raw table in comma separated value (CSV) format. The summary table contains the 15 traits listed in Table 2, and is calculated for each image in the input image folder. The raw table contains the individual RHL and RHA measurements from each bin in the sliding window for each input image.

Users can quickly view the summary information displaying RHL and RHA profile for each image via the --plot-summary flag. To visualize the segmentation and transformation of the input image, --plot-segmentation saves the generated binary mask of each image. To view how the straightening was performed, --plot-transformation saves a graphical representation of the root warping. To operate pyRootHair, users only need to provide three required arguments: **-i/--input** specifies the filepath/directory to the input image folder. **-b/--batch-id** specifies the sub-folder name associated with the current run, which is stored in the output directory. **-o/--output** specifies a filepath/directory to store the output data and plots.

Since varying root length in the input image affects how summary traits (e.g. Mean RHL, total RHA) are calculated, we offer the argument --length-cutoff, which allows users to specify a length cutoff for input roots (measured in milimetres from the root tip). --length-cutoff standardises trait calculation for all images in the input batch based on the value provided. For example, --length-cutoff 10 will return summary and raw tables of only the first 10 millimetres of the root for all images in the batch. This mitigates a significant portion of bias introduced with varying root length, enabling standardisation of traits for more detailed analysis.

A detailed breakdown of all the arguments and flags is available on the following Github repository https://github.com/iantsang779/pyRootHair.

### 8.7 Dependencies

pyRootHair was written in the Python programming language (v3.12.7) and developed on the Crop Diversity HPC, running the Debian 12 Bookworm operating system (Percival-Alwyn et al. 2024). Scikit-image (v0.24.0) was used to carry out most image processing functionalities. Numerical calculations were carried out using the numpy (v2.0.2) and scipy (v1.14.1) libraries. Data tables were constructed using pandas (v2.2.3) and all plots and figures were created using matplotlib (v3.9.3). LOWESS regression lines were computed using statsmodels (v0.14.4). Scikit-learn (v.1.5.2) was used for quality control of segmented images. nnU-Netv2 (v2.5.1) was used to create the image segmentation model with PyTorch (v.2.5.1) and CUDA (v.12.6).

### 8.8 Installation

Extensive documentation and source code is available from the github repository: https://github.com/iantsang779/pyRootHair. pyRootHair is available on pip for installation: https://pypi.org/project/pyRootHair/

## 9 Availability of Source Code and Requirements

All code used to generate figures for this paper has been made available in the following Jupyter Note-book: https://github.com/iantsang779/pyRootHair/blob/main/paper_data/pyRootHair.ipynb. The source data used to generate figures is available here: https://github.com/iantsang779/pyRootHair/tree/main/paper_data.

- Project name: pyRootHair
- Project home page: https://github.com/iantsang779/pyRootHair
- Operating system(s): Linux, MacOS, Windows
- Programming language: Python
- License: MIT License

## 10.2 List of Abbreviations

AI: Artifical Intelligence
CNN: Convolutional Neural Network
GPU: Graphical Processing Unit
GWAS: Genome Wide Association Studies
HPC: High Performance Computer
LOWESS: Locally Weighted Scatterplot Smoothing
RHL: Root Hair Length
RHA: Root Hair Area
RHD: Root Hair Density
VRAM: Video Random Access Memory

## 10.4 Author Contributions

I.T - Conceptualization, Data curation, Formal analysis, Methodology, Software, Validation, Visualization, Writing - original draft, Writing - review & editing. L.P.A - Validation, Writing - review & editing. S.R, J.C, F.L, J.A.A - Supervision, Writing - review & editing.

## 10.5 Acknowledgements

The authors would like to thank Marjorie Guichard for providing arabidopsis, medicago and brachypodium images for training, and Nina Foreman for providing rice seed stocks. We also thank Mike Pound and Feng Chen for their invaluable advice on CNNs. The authors acknowledge Research Computing at the James Hutton Institute for providing computational resources and technical support for the “UK’s Crop Diversity Bioinformatics HPC” (BBSRC grants BB/S019669/1 and BB/X019683/1), use of which has contributed to the results reported within this paper.

## 13 Supplementary Figures

**Supplementary Figure 1:**
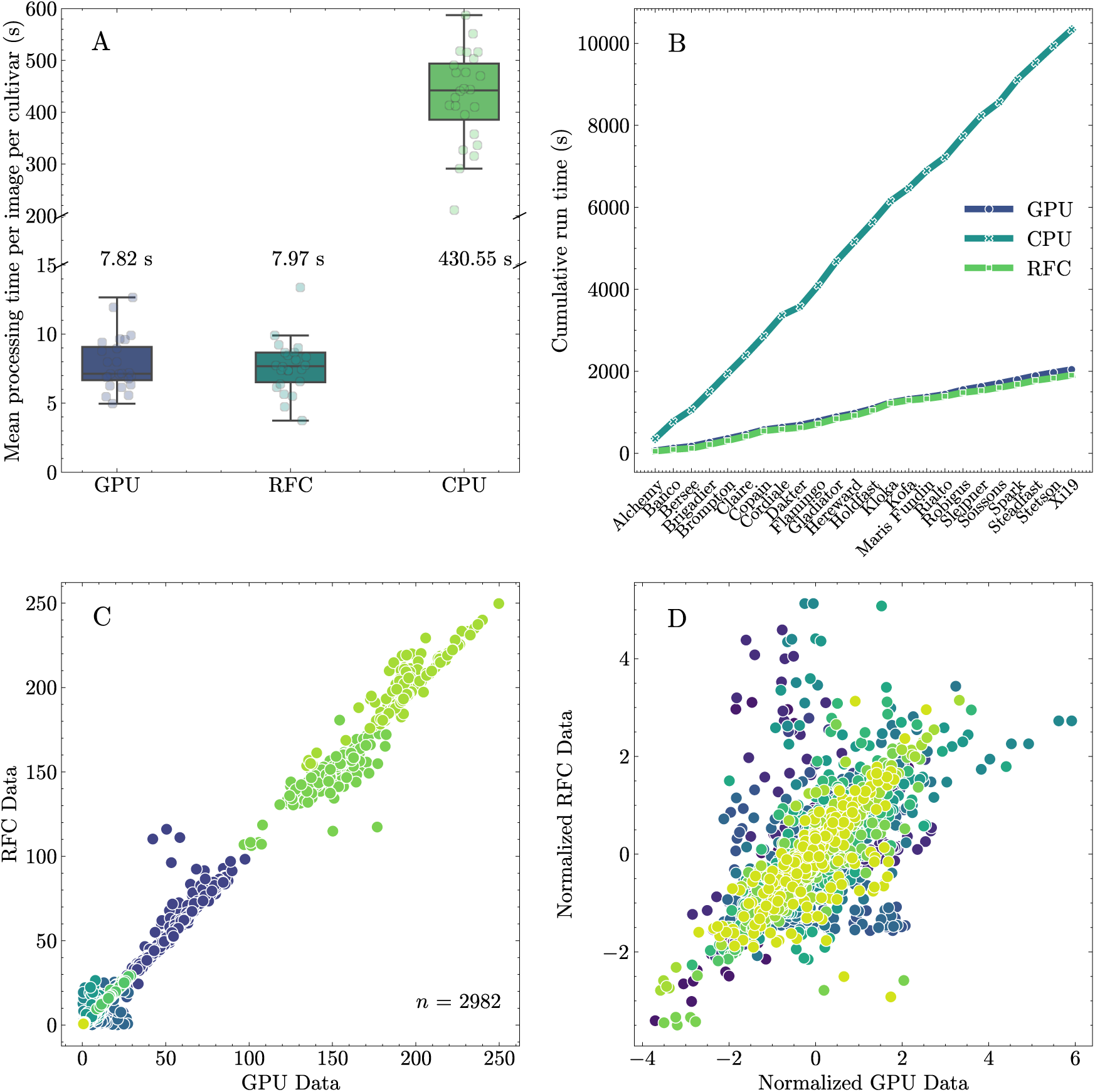
Comparing processing times between different pyRootHair pipeline configurations. A) Boxplot illustrating the mean processing duration in seconds (including inference) for each image within each cultivar input folder. Mean runtime per image annotated above/below each box. CPU pipeline utilized the same CNN deployed in the main (GPU) pipeline to perform inference without a GPU. B) Cumulative run time for all 252 images used in this study across all three pipeline options. (C) Summary data and (D) mean normalized summary data calculated for all images using the default pipeline (GPU) and the random forest classifier (RFC) pipeline. The GPU pipeline was run using an Nvidia L40S GPU with 8 GB VRAM. The RFC pipeline was run on an HPC compute node with 20 GB RAM using a RFC model trained on a single image. CNN: convolutional neural network. GPU: Graphical Processing Unit. RFC: Random Forest Classifier. CPU: Central Processing Unit, HPC: High Performance Computer

**Supplementary Figure 2:**
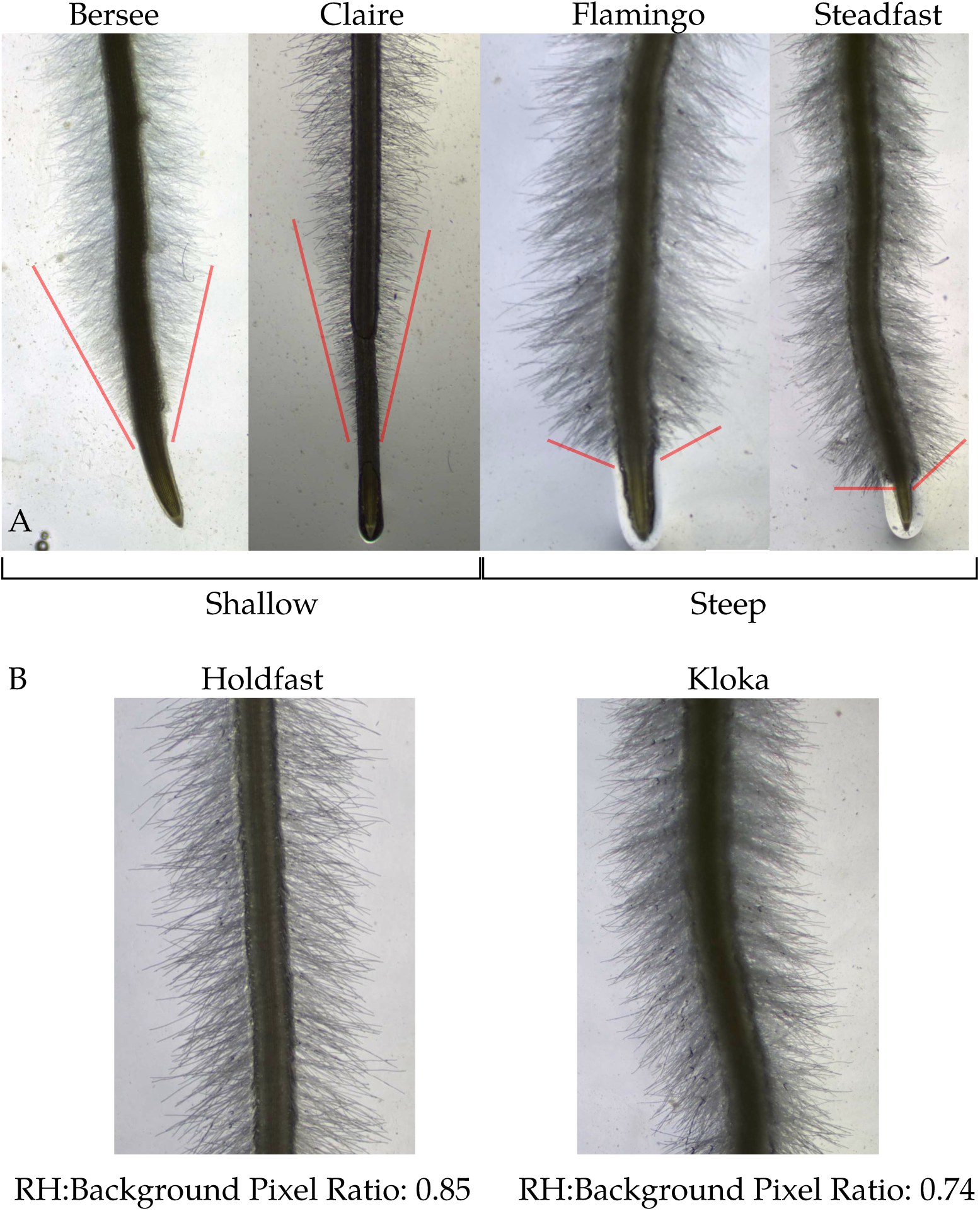
Highlighting variation in root hairs around root tips. A) Representative images of root tips of wheat cultivars with ‘shallow’ (Bersee, Claire) and ‘steep’ (Flamingo, Steadfast) root hair profiles as identified in Figure 5. Red lines illustrate the difference in gradient of the root hair profile at root tip. B) A visual representation between cultivars with a higher (Holdfast) and lower (Kloka) RH:Background Pixel Ratio. Low values indicate increased root hair area (RHA), while high values indicate low RHA.

**Supplementary Figure 3:**
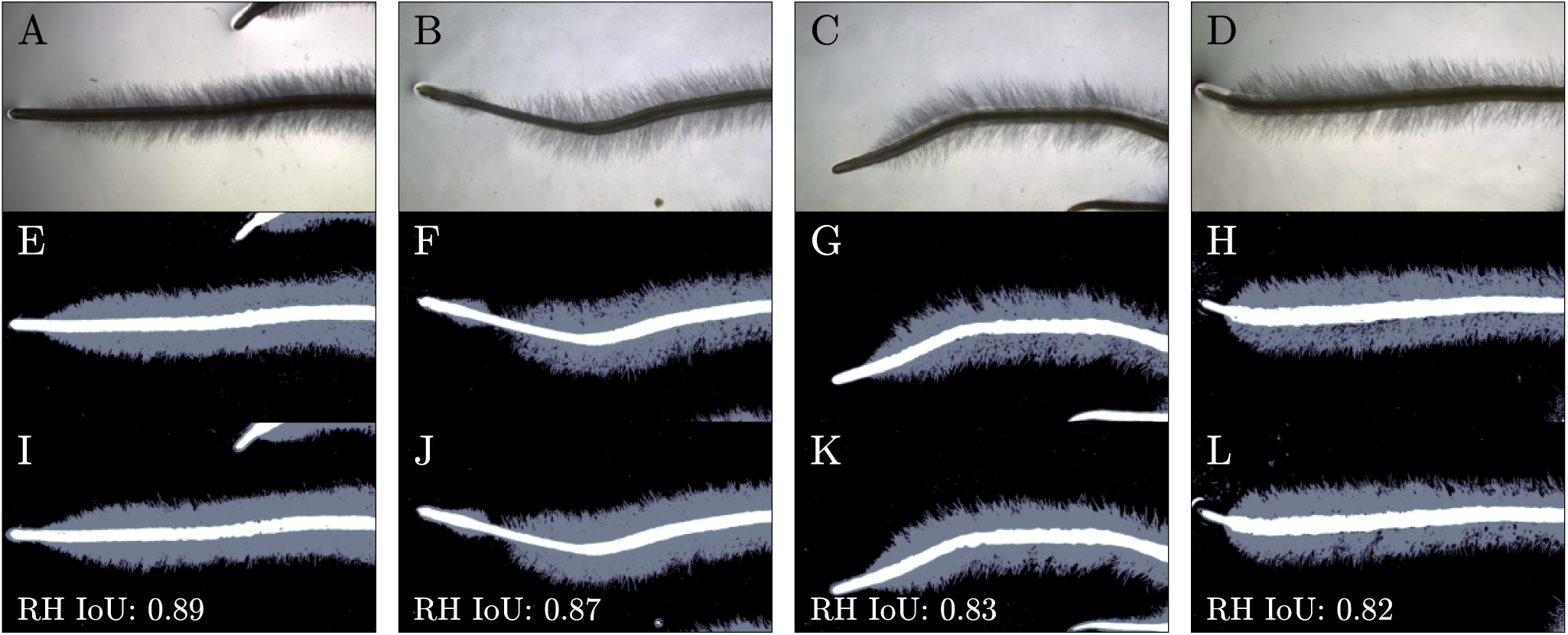
Validation of Convolutional Neural Network (CNN) segmentation on four selected images. A-D) Raw Images, E-H) Manually annotated binary masks of the above images using ilastik (Berg et al. 2019), I-L) Predicted binary masks generated from the CNN. Intersection over union (IoU) scores for the root hair masks are shown at the bottom. IoU scores illustrate proportion of overlapping pixels in the manually annotated (E-H, grey) and the predicted root hair masks (I-L, grey), divided by the total area of root hair masks minus the overlapping region (intersection).

